# Exome Sequencing and Prediction of Long-Term Kidney Allograft Function

**DOI:** 10.1101/015651

**Authors:** L. Mesnard, T. Muthukumar, M. Burbach, C. Li, H. Shang, D. Dadhania, J. R Lee, V. K. Sharma, J. Xiang, C. Suberbielle, M. Carmagnat, N. Ouali, E. Rondeau, J. J. Friedewald, M. M. Abecassis, M. Suthanthiran, F. Campagne

**Affiliations:** The HRH Prince Alwaleed Bin Talal Bin Abdulaziz Alsaud Institute for Computational Biomedicine, Weill Cornell Medical College, New York, NY, United States of America; Department of Physiology and Biophysics, The Weill Cornell Medical College, New York, NY, United States of America; Division of Nephrology and Hypertension, Weill Cornell Medical College, New York, NY, United States of America; Department of Transplantation Medicine, New York Presbyterian Hospital, New York, NY, United States of America; Genomics Core Facility Weill Cornell Medical College; Northwestern University Feinberg School of Medicine, Chicago, IL, United States of America; Comprehensive Transplant Center, Northwestern University Feinberg School of Medicine, Chicago, IL, United States of America; INSERM UMR1155 et Service des Urgences Néphrologiques et Transplantation Rénale, APHP, Hôpital Tenon, 75020 Paris, France; Laboratoire d’histocompatibilité Hôpital Saint Louis APHP, Paris, France

## Abstract

**Abstract:** Current strategies to improve graft outcome following kidney transplantation consider information at the HLA loci. Here, we used exome sequencing of DNA from ABO compatible kidney graft recipients and their living donors to determine recipient and donor mismatches at the amino acid level over entire exomes. We estimated the number of amino acid mismatches in transmembrane proteins, more likely to be seen as foreign by the recipient’s immune system, and designated this tally as the allogenomics mismatch score (AMS). The AMS can be measured prior to transplantation with DNA for potential donor and recipient pairs. We examined the degree of relationship between the AMS and post-transplantation kidney allograft function by linear regression. In a discovery cohort, we found a significant inverse correlation between the AMS and kidney graft function at 36 months post-transplantation (n=10 recipient/donor pairs; 20 exomes) (r^2^>=0.57, P<0.05). The predictive ability of the AMS persists when the score is restricted to regions outside of the HLA loci. This relationship was validated using an independent cohort of 24 recipient donor pairs (n=48 exomes) (r^2^>=0.39, P<0.005). In an additional cohort of living and mostly intra-familial recipient/donor pairs (n=19, 38 exomes), we validated the association after controlling for donor age at time of transplantation. Finally, a model that controls for donor age, HLA mismatches and time post-transplantation yields a consistent AMS effect across these three independent cohorts (P<0.05). Taken together, these results show that the AMS is a strong predictor of long-term graft function in kidney transplant recipients.

**One Sentence Summary:** Prediction of long-term kidney graft function with exome sequencing

## Introduction

Survival of patients afflicted with End Stage Renal Disease (ESRD) is superior following kidney transplantation compared to dialysis therapy. The short-term outcomes of kidney grafts have steadily improved since the early transplants (performed in the 1960s) with refinements in immunosuppressive regimens, use of DNA-based HLA typing, and better infection prophylaxis *(1–3)*. Despite these advances, data collected across the USA and Europe show that 40-50% of kidney allografts fail within ten years of transplantation *(4)*. This observation strongly suggests that as yet uncharacterized factors, including genomic loci, may adversely impact long-term post-transplantation outcomes.

Observational studies have demonstrated the importance of matching for the HLA-determined proteins on kidney graft outcome. Therefore, in many countries, including the USA, donor kidney allocation algorithms includes consideration of HLA matching. With widespread incorporation of HLA matching in organ allocation decisions, it has become clearer that HLA mismatching represents an important risk factor for kidney allograft failure but fails to fully account for the invariable decline in graft function and failure in a large number of cases over time. Indeed, only a 15% survival difference exist at 10 years post transplantation between the fully matched kidneys and the kidneys mismatched for both alleles at the HLA-A, B and DR loci *(5)*. These observations suggest that mismatches at non-HLA loci in the genome could influence long-term graft outcomes. The current clinical practice of prescribing life-long immunosuppressive therapy to recipients of fully HLA matched donor kidneys, but not to recipients of monozygotic identical twin kidneys, also suggests a role for non-HLA related genomic factors on graft outcome.

While tests of allelic frequencies are a hallmark of genetic research, transplantation has none of the Mendelian characteristics for which genetic tests have been developed. Therefore, the assumption of the Mendelian transmission model seems inadequate to develop predictors of graft function following transplantation. Indeed, previous attempts at using this methodology have identified small genotype effects on graft function in cohorts of hundreds of transplant patients, but often could not be replicated in independent cohorts (reviewed in *(6)*).

In this work, we present a new method to estimate the genomic compatibility between the organ graft recipient and donor. This approach, designated as allogenomics in this communication, considers the entire coding sequence of both recipient and donor genomes, as determined by exome sequencing. The allogenomics concept makes it possible to estimate a quantitative compatibility score between the genomes of a recipient and potential donor and is calculated from genotypes and genome annotations available before transplantation. The allogenomics approach does not assume a Mendelian inheritance model but integrates the unique features of transplantation such as the existence of two genomes in a single individual and the recipient’s immune system mounting an immune response directed at antigens displayed by the donor kidney. In this report, we show that this new concept helps predict long-term kidney transplant function from the genomic information available prior to transplantation.

## Results

### The allogenomic concept and the allogenomics mismatch score (AMS)

The allogenomics concept is the hypothesis that interrogation of the coding regions of the entire genome for both the organ recipient and organ donor DNA can identify the number of incompatible amino-acids (recognized as non-self by the recipient) that inversely correlates with long-term graft function post transplantation. **Fig. 1A** is a schematic illustration of the allogenomics concept. Because human autosomes have two copies of each gene, we consider two possible alleles in each genome of a transplant pair. To this end, we estimate allogenomics score contributions between zero and two, depending on the number of different amino acids that the donor genome encodes for at a given protein position. **Fig. 1B** shows the possible allogenomics score contributions when the amino acids in question are either an alanine, or a phenylalanine or an aspartate amino acid. The allogenomics mismatch score (AMS) is a sum of amino acid mismatch contributions. Each contribution represents an allele coding for a protein epitope that the donor organ may express and that the recipient immune system could recognize as non-self (see Equation 1 and 2 in **Fig. 1C** and Materials and Methods and full description in the supplementary appendix).

**Figure 1.**
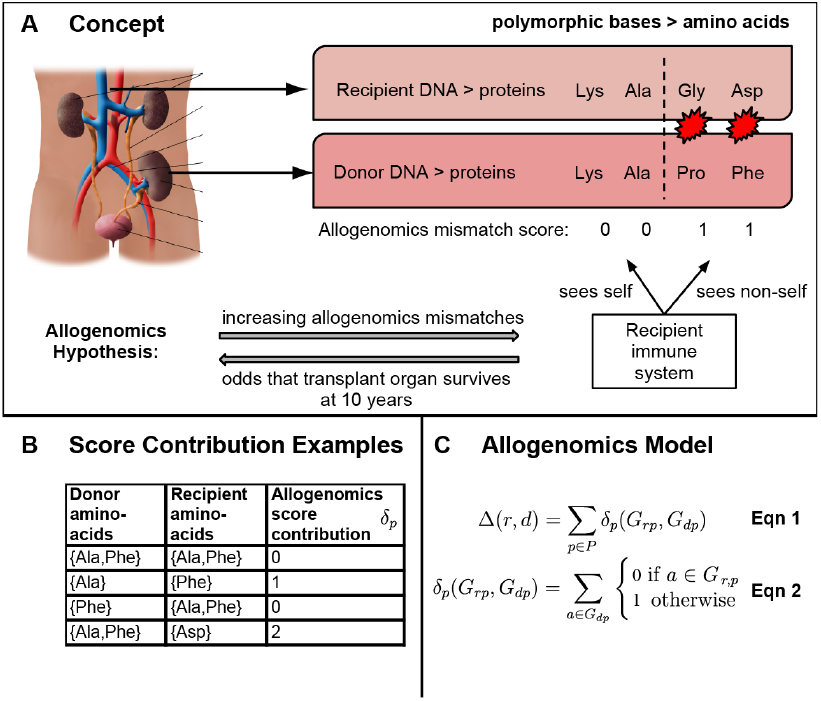
Recipient/Donor incompatibility quantified by exome sequencing and calculation of allogenomics mismatch score (AMS). **(A)** Hypothesis: Post-transplantation kidney graft function is associated with the number of amino acids coded by the donor genome that the recipient’s immune system could recognize as non-self. **(B)** Examples of donor/recipient amino711 acid mismatches at one protein position, and resulting contributions to the allogenomics mismatch score. The allogenomics mismatch score is calculated by summing contributions over a set of genomic polymorphisms (see Methods for details). **(C)** Equations for the allogenomics model. Score contributions are summed across all genomic positions of interest (set *P*) to yield the allogenomics score Δ(*r,d*). *G_r,p_*: genotype of recipient *r* at genomic site/position *p. G_d,p_*: genotype of donor *d* at site *p*. Alleles of a genotype are denoted with the letter *a*.

We have developed and implemented a computational approach to estimate the allogenomics mismatch score from genotypes derived for pairs of recipient and donor genomes. (See Materials and Methods for a detailed description of this approach and its software implementation, the allogenomics scoring tool, available at http://allogenomics.campagnelab.org.) Our approach is designed to consider the entire set of protein positions measured by a genotyping assay, or restrict the analysis to a subset of positions *P* in the genome. In this study, we focus on the subset of genomic sites P that encode for amino acids in transmembrane proteins.

### The AMS correlates with post-transplantation graft function in living donor-kidney recipient pairs

In order to test the allogenomics hypothesis, we isolated DNA from 10 kidney graft recipients and their living donors (Discovery Cohort), performed whole exome sequencing and analyzed genotype data for these recipient/donor genome pairs (10 pairs, 20 exomes). These patients were a subset of patients enrolled in a multicenter Clinical Trial in Organ Transplantation-04 (CTOT- 04) study of urinary cell mRNA profiling, from whom tissue/cells were collected for future mechanistic studies *(7)*. **Table 1** provides demographic and clinical information about the patients included in the Discovery Cohort. Exome data were obtained with the Illumina TrueSeq exome enrichment kit v3, covering 62Mb of the human genome. Primary sequence data analyses were conducted with GobyWeb *(8)* (data and analysis management), Last *(9)* (alignment to the genome) and Goby *(10)* (genotype calls).

**Table 1.** Characteristics of Kidney transplant recipients and their donors. In bold, characteristics that differ between the French *and* Validation cohorts (*P<0.05, two tailed t-test).

| Characteristic | Discovery cohort | Validation cohort | French Cohort |
| --- | --- | --- | --- |
| Number of Transplant Pairs with living donors | 10/10 | 24/24 | 19/19 |
| Allogeneomics mismatch score AMS(SD)[range] | 1335(304)[994-2033] | 1094(259)[700-1630] | 560(147)[349-811] |
| Clinical factors |  |  |  |
| Age |  |  |  |
| Donor (SD) | 41 (13) | 46 (10) | 44(16) |
| Recipient (SD) | 48 (10) | 51 (13) | 38(15) |
| Living Donor type |  |  |  |
| Living related N (AMS) [SD]) | 4 (1116 [143]) | 13 (939 [218]) | **15(503[108])\*** |
| Living unrelated N (AMS) [SD]) | 6 (1481 [300]) | 11(1277 [170]) | **4(769[41])\*** |
| Donor sex |  |  |  |
| Male (%) | 2 (20%) | 8 (33%) | 6(32%) |
| Female (%) | 8 (80%) | 16 (67%) | 13(68%) |
| Donor Race |  |  |  |
| Black (%) | 4(40%) | 5 (21%) | 2(10%) |
| Non-Black (%) | 6(60%) | 19 (79%) | 17(90%) |
| Recipient sex |  |  |  |
| Male (%) | 9 (90%) | 13 (54%) | 13 (53%) |
| Female (%) | 1 (10%) | 11 (46%) | 13 (47%) |
| Recipient Race |  |  |  |
| Black (%) | 4 (40%) | 7 (29%) | 2 (10%) |
| Non-Black (%) | 6 (60%) | 17 (71%) | 17 (90%) |
| Number of HLA mismatches ABDR (SD) | 3.9 (1.91) | 3.5 (1.89) | 2.5 (1.68)* |
| Functional Factors |  |  |  |
| Number of Patients at 12 months | 10 | 24 | 17 |
| Serum creatinine level at 12 months mg/dL (SD) | 1.51 (0.35) | 1.45 (0.41) | 1.29 (0.41) |
| eGFR at 12 months ml/min/1.73m<sup>2</sup> (SD) | 54.3(10) | 54.3 (16.3) | **61.8 (18.9)\*** |
| Number of Patients at 24 months | 9 | 23 | 19 |
| Serum creatinine level at 24 months mg/dL (SD) | 1.36 (0.19) | 1.45 (0.49) | 1.26 (0.3) |
| eGFR at 24 months ml/min/1.73m<sup>2</sup> (SD) | 59 (7.7) | 54.85 (15.7) | **59.3 (14.5)\*** |
| Number of Patients at 36 months | 8 | 22 | 19 |
| Serum creatinine level at 36 months mg/dL(SD) | 1.62 (0.50) | 1.38 (0.40) | 1.35 (0.45) |
| eGFR at 36 months ml/min/1.73m<sup>2</sup> (SD) | 53.4 (15) | 55.3 (15.9) | 56.3 (16.4) |
| Number of Patients at 48 months | 0 | 16 | 16 |
| Serum creatinine level at 48 months mg/dL(SD) | - | 1.34 (0.43) | 1.40 (0.56) |
| eGFR at 48 months ml/min/1.73m<sup>2</sup> months (SD) | - | 57.4 (16.4) | 55.7 (18.2) |
| Patients with an Acute Cellular rejection episode in the first year of transplantation, N (%) | 3 (30%) | 5 (20%) | 2 (10%) |
| Immunosuppression |  |  |  |
| Calcineurin Inhibitors, n (%) | 9 (90%) | 24 (100%) | 19 (100%) |
| Corticosteroids, n (%) | 0 (0%) | 5 (21%) | **17 (90%)\*** |

Kidney graft function is a continuous phenotype and is clinically evaluated by measuring serum creatinine levels or using estimated glomerular filtration rate (eGFR)*(11)*. In this study, kidney graft function was evaluated at months 12, 24, 36 and/or 48 following transplantation using serum creatinine levels and eGFR, calculated using the 2011 MDRD *(12)* formula. We examined whether the allogenomics mismatch score is associated with post-transplantation allograft function.

We found positive linear associations between the allogenomics mismatch score and serum creatinine level at 36 months post transplantation (r^2^ adj.=0.78, P=0.002, n=10, at 36 months) but not at 12 or 24 months following kidney transplantation (**Fig. 2A, B, C**). We also found a negative linear relationship between the score and eGFR at 36 months post transplantation (r^2^ adj.=0.57, P=0.02) but not at 12 or 24 months following kidney transplantation (**Fig. 2D, E, F**). These findings suggest that in the discovery cohort the allogenomics score is predictive of long-term graft function, but perhaps not short-term function.

**Figure 2.**
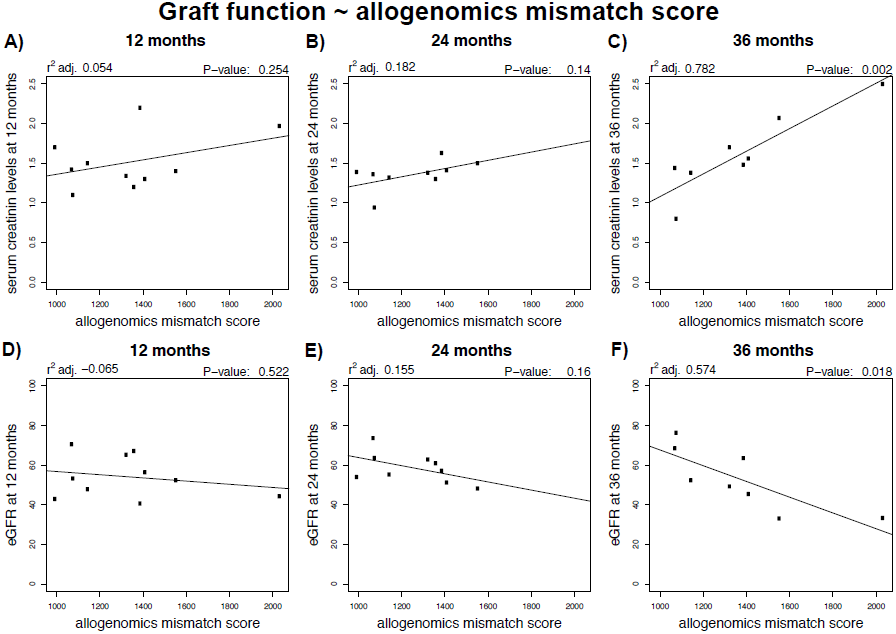
Relationship between the allogenomics mismatch score (AMS) and kidney graft function at 12, 24 or 36 months following transplantation in the Discovery cohort. DNA was isolated from 10 pairs of kidney graft recipients and their living kidney donors (Discovery set). Whole exome sequencing of the donor genomes and recipient genomes was performed and the sequencing information was used to calculate allogenomics mismatch scores based on amino acid mismatches in transmembrane proteins. The panels depict the relationship between the allogenomics mismatch scores and serum creatinine levels at 12, 24 and 36 months post transplantation (Panels A, B and C, respectively) and the relationship between the allogenomics mismatch scores and estimated glomerular filtration rate at 12, 24 and 36 months post transplantation (Panels D, E and F, respectively). Both serum creatinine levels and eGFR correlate in a time-dependent fashion with the allogenomics mismatch score with the strongest correlations being observed at 36 months post-transplantation.

### The AMS correlates with graft function in a second, independent cohort of non-related living kidney donor/recipient pairs

We sought to validate the observation that the AMS is associated with post-transplantation kidney graft function by testing the association in an independent cohort of kidney transplant patients. To this end, we sequenced DNA collected from 24 additional kidney donor/recipient pairs (see **Table 1** for information about subjects included in the Validation cohort). DNA sequencing was performed using the Agilent Haloplex assay covering 37Mb of the coding sequence of the human genome. We called the genotypes and estimated the AMS as described for the discovery cohort (see Methods). **Fig. 3** shows that, as observed with the Discovery cohort, the AMS correlates progressively better with kidney graft function at longer times following transplantation. At 36 months post transplantation, a small to moderate positive association was observed between the allogenomics mismatch score and the serum creatinine level (r^2^ adj. 0.139, P=0.049) (**Fig. 3C**) and eGFR (r^2^ adj. 0.078, P=0.11) (**Fig. 3G**). The association between the score and graft function was stronger and reached significance at 48 months post transplantation for both creatinine level (r^2^ adj. 0.394, P<0.01) (**Fig. 3D**), and eGFR (r^2^ adj. 0.284, P=0.02) (**Fig. 3H**), further validating the association in the Validation cohort. In order to test whether models trained on one cohort would generalize to another similar cohort, we trained models on the Discovery cohort and used the fixed model to predict graft function in the Validation cohort. **Fig. S1** shows that such a fixed model does generalize when presented with new recipient-donor pairs, and also exhibited better fit to the longer 48-month time-point compared to the earlier time point (**Fig. S1B** vs. **Fig. S1C**). Similarly, models trained on the Validation cohort generalize to the Discovery cohort (**Fig. S2**). These results establish that the parameters of the models are stable, despite the relatively small numbers of kidney recipient-donor pairs included in the Discovery and Validation cohorts.

**Figure 3.**
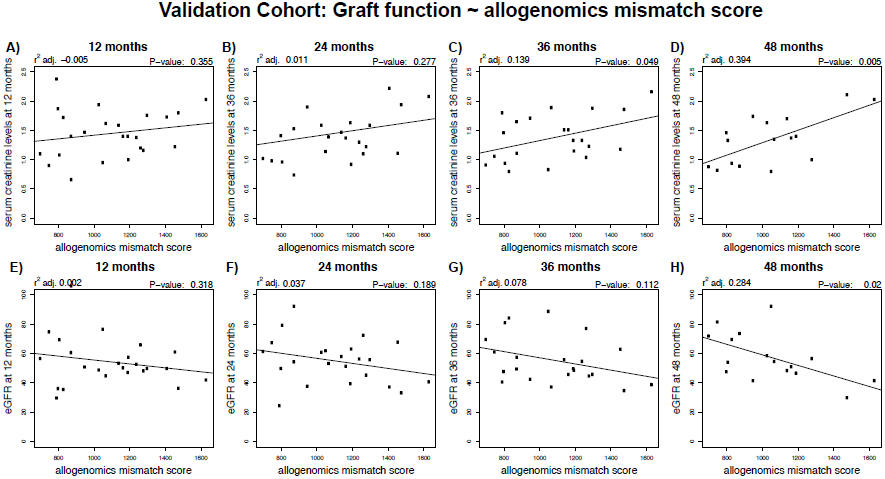
Relationships between allogenomics mismatch scores (AMS) and kidney graft function at 12, 24, 36 or 48 months post transplantation in the Validation Cohort. DNA was isolated from 24 pairs of kidney graft recipients and their living kidney donors (Validation set). Whole exome sequencing of the donor and recipient genomes was performed and the sequencing information was used to calculate allogenomics mismatch scores based on amino acid mismatches in transmembrane proteins. The relationships between allogenomics mismatch scores and serum creatinine levels at 12, 24, 36 and 48 months post transplantation (Panels A, B, C, and D respectively) are shown. In panels E, F, G and H, the relationships between the allogenomics mismatch scores and estimated glomerular filtration rate at 12, 24, 36 and 48 months post transplantation are shown. Both serum creatinine levels and eGFR correlate in a time dependent fashion with the allogenomics mismatch score with the strongest correlations being observed at 48 months post-transplantation.

### The AMS weakly correlates with graft function in a third cohort

In order to further test the strength of the relationship between AMS and graft function we applied the approach to a third independent cohort. To this end, we assembled a new cohort composed mostly of living related kidney donor pairs. We used a peculiarity of the French transplant system, directed by the French national agency for organ procurement transplant, which until recently did not allow living non-related kidney transplantation, to select 19 additional pairs from one French center. Demographic and clinical data for this cohort are shown in **Table 1**. As expected in this situation, the range of the AMS is lower than in the other two cohorts, ranging from 349 to 811 in the well-matched cohort (mean 559 ± 147). This range was 700 to 1630 (mean 1094 ± 259) in the validation cohort and 994 to 2033 in the discovery cohort (mean 1335 ± 304). These data suggest that the effective range of variation of the AMS is approximately 300-2000 (1700 AMS units).

In this third cohort, we did not find the simple association between the AMS and graft function that we observed in the first two cohorts. However, after correction for the strong effect of donor age, we observe a trend in the same direction as the association observed in the first two cohorts at 24, 36 and 48 months when predicting eGFR (e.g., AMS effect estimate=-0.021798 at 36 months). The three cohorts combined yield a statistically significant effect of AMS on graft function at 36 months (n=48 pairs, AMS effect estimate=-0.015808, P<0.01) and 60 months (n=13 pairs, AMS effect estimate=-0.04479, P<0.01), and borderline at 48 months (n=32 pairs, AMS effect estimate=-0.016685, P=0.07).

### Analysis of the relationship between the AMS and the number of HLA mismatches

HLA mismatches are well-described risk factors for kidney graft failure. Therefore given data obtained in the first two cohorts, we next focused our attention on the association between AMS and number of ABDR HLA mismatches. **Fig. 4** presents an analysis where we combined the Discovery and Validation cohorts (32 transplant kidney recipient-donor pairs) and compared the AMS to the number of mismatches at the HLA-A, B and DR loci. We find that the AMS was moderately correlated with the number of mismatch at the HLA loci (**Fig. 4A**, r^2^ adj.=0.35, P<0.001). However, the number of HLA mismatches correlates poorly with an AMS estimated from exome data when restricting the sites to the HLA A, B and DR loci (**Fig. 4B**, r^2^ adj.=0.09, P=0.047). Furthermore, the AMS estimated outside of the HLA A, B and DR loci still associates significantly with serum creatinine levels (**Fig. 4C**, r^2^ adj.=0.36, P<0.001 and eGFR (**Fig. 4D**, r^2^ adj.=0.18, P=0.011). These data indicate that the ability of the AMS to predict future graft function is mostly independent of the HLA loci.

**Figure 4.**
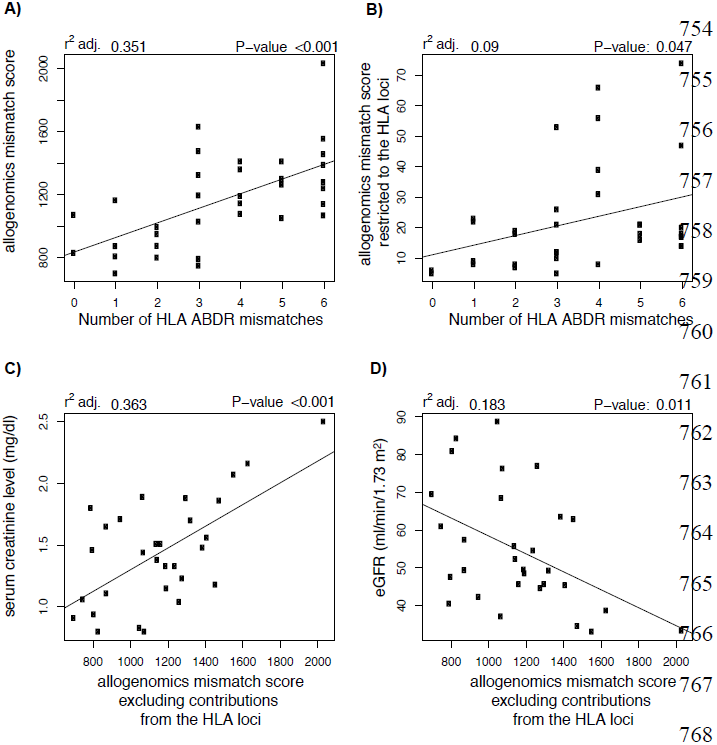
Relationships between allogenomics mismatch scores and the HLA loci. We combined the Discovery and Validation cohorts to examine the relation of the allogenomics mismatch score to the HLA-A, B and DR mismatch between the recipient and the kidney donor. Allogenomics mismatch scores, either calculated over all transmembrane proteins (Panel A), or restricted to the HLA A, B DR loci (Panel B) correlate with the number of mismatches in the A,B, and DR HLA loci for the complete cohort. These correlations however do not explain the association with post-transplantation graft function because when the HLA loci (A/B/ DR/DQ) are excluded from the sites included in the calculation of the allogenomics mismatch score, a significant correlation is still observed between the allogenomics mismatch score and serum creatinine level (Panel C) and eGFR (Panel D) at 36 months post transplantation (similar results are obtained when excluding the A/B/C/DR/DQ/DP HLA loci, data not shown).

### Final predictive models

We fit models across the three combined cohorts to yield final models with fixed parameters:

36-month eGFR = 106.311073 -0.015808 AMS -0.749321*Donor_Age
48-month eGFR = 108.553088 -0.016685*AMS -0.798154*Donor_Age
60-month eGFR = 138.60095 -0.04479*AMS -1.14063 *Donor_Age

These equations and parameters are provided to enable testing these models on independent cohorts of living transplant pairs genotyped with exome sequencing. Fit parameter values were estimated with MetaR (train linear model statement, version 1.5.0). We note that the fixed parameters of this model can be sensitive to the exact analysis pipeline used to align reads to the genome and to call genotypes and that an objective test of this model would follow the analysis protocols used to analyze data for this report.

In the models presented so far, we have considered the prediction of graft function separately at different time points. An alternative analysis would consider time since transplantation as another factor that can influence graft function. This is particularly useful when studying cohorts where graft function was assessed at several distinct time points (e.g., in the French cohort, clinical data describes graft function from 1 to 96 months post transplantation, but few time points have observations for all recipients). To implement this alternative analysis, we fit a mixed linear model of the form eGFR ~ donor age at time of transplant + AMS + T + (1|P), where T is the time post-transplantation, measured in months, and (1|P) a random effect which models separate model intercepts for each donor/recipient pairs. To determine the effect of AMS on eGFR, we compare the fit of models that do or do not include AMS. We find that the effect of AMS is significant (P=0.0042, χ2= 8.1919, d.f.=1). A similar result is obtained if HLA is used in covariate in the model (i.e., eGFR ~ donor age at time of transplant + AMS + T + HLA + (1|P), comparing model with AMS or without, P= 0.038, χ2= 4.284, d.f.=1). In contrast, models that include AMS, but do or do not include the number of ABDR HLA mismatches fit the data equally well (P= 0.60, χ2= 0.2737, d.f.=1), confirming that the effect of AMS is independent of the number of HLA mismatches.

Table 2 presents confidence intervals for the parameters of the full model (eGFR ~ donor age at time of transplant + AMS + T + HLA + (1|P)) as well as the effective range of the model predictors. The table shows the expected impact of each predictor on eGFR when this predictor is varied over its range, assuming all other predictors are kept constant. For instance, we assume that donor age at time of transplant varies from 20 years old to 80 years old (range: 60). Across this range, eGFR will decrease by an estimated 28 units as the donor gets older. The AMS effect has an effective range of 1,700 and the corresponding eGFR decrease is 19 units.

**Table 2.** Estimated Model Parameters, 95% Confidence Intervals and Expected Impact on eGFR.

| Model Coefficient | Estimate |  |  | Effective Range | Impact on eGFR |  |  | Note |
| --- | --- | --- | --- | --- | --- | --- | --- | --- |
|  | Fit | 2.50% | 97.50% |  | Fit | 2.50% | 97.50% |  |
| Time post transplantation (in months) | -0.2439 | -0.32348868 | -0.1644768 | 480 | -117.072 | -155.2745664 | -78.948864 | * |
| Donor age at transplant (in years) | -0.46843 | -0.78098518 | -0.15179 | 60 | -28.1058 | -46.8591108 | -9.1074 | * |
| AMS | -0.01141 | -0.02217172 | -0.000627551 | 1700 | -19.397 | -37.691924 | -1.06683704 | * |
| HLA_ABDR_mismatches | -0.57055 | -2.73627999 | 1.606998 | 6 | -3.4233 | -16.41767994 | 9.641988 | n.s. |

### Impact of genotyping platform on the estimation of the AMS

We studied the impact of the genotyping platform on the estimation of the AMS (**Fig. S3**). Large cohorts of matched recipient and donor DNA are being assembled and genotyped with SNP chip array technology such as the Illumina 660W bead array platform *(13)*. We asked whether such platforms would be appropriate to validate the allogenomics model in large cohorts. **Fig. S3A** documents the number of sites that contribute to the allogenomics score on each platform. **Fig. S3B** indicates that the exome assay captures many more sites with rare polymorphisms (minor allele frequency <5%) than the GWAS array platform. This is expected because exome assays directly sequence an individual DNA, while GWAS platforms are designed with a fixed set of polymorphisms and will not include many of the rare polymorphisms any given individual may carry. **Fig. S3C** compares the correlations measured with the exome assay or that could have been obtained if we had measured the allogenomics mismatch score with the Illumina 660W assay. The weak correlations obtained suggest that GWAS platforms are not ideal for future tests of the allogenomics model.

## Discussion

Several Donor/Recipient matching factors have been identified prior to this study as important for transplantation. For instance, blood group compatibility is a prerequisite unless pre-conditioning of the recipient is undertaken to facilitate blood group incompatible kidney transplantation. While HLA compatibility is a necessary requirement for successful bone marrow transplants, full HLA compatibility is not an absolute prerequisite for all types of transplantations as indicated by the thousands of solid organ transplants performed yearly despite lack of full matching between the donor and recipient at the HLA-A, B and DR loci. In view of better patient survival following transplantation compared to dialysis, kidney transplants are routinely performed with varying degrees of HLA-mismatches, including HLA mismatches for all HLA-class I and II antigens. Although, graft outcomes improve with better HLA-matching, excellent long-term graft outcomes with stable graft function have been observed in patients with full ABDR HLA mismatches. The success of these transplants clearly suggests that factors other than HLA compatibility may influence the long-term clinical outcome of kidney allografts.

Case-control designs are appropriate when studying phenotypes that are expected to be associated with genotypes that follow a Mendelian inheritance mechanism. We reason that transplant patients are not ideal subjects for this experimental design. Indeed, patients who received a kidney transplant have two genomes in their body: their germline DNA, and the DNA of the donor. In cases when the transplanted kidney was from an unrelated donor (e.g., organs from deceased donors), it is clear that a Mendelian genetic transmission mechanism is not at play. Importantly, even in cases where the donor is one of the parents of the transplant recipient (familial, living related transplant), the genome of the parent will break the assumptions of Mendelian inheritance. Because the transplant recipient has two genomes Mendelian inheritance. Because the transplant recipient has two genomes after transplant, it is not appropriate to assume that genomic markers can be identified when assuming a Mendelian inheritance process. Yet, this assumption has been made in most of the transplantation genomic studies published to date.

The allogenomics concept that we present in this manuscript postulates a different mechanism for the development of the immune response in the transplant recipient: immunological and biophysical principles strongly suggest that alleles present in the donor genome, but not in the recipient genome, will have the potential to produce epitopes that the recipient immune system will recognize as non-self. This reasoning explains why the allogenomics score is not equivalent to the genetic measures of allele sharing distance that have been used to perform genetic clustering of individuals *(14)*.

Results presented in this manuscript suggest that allogenomic mismatches in proteins expressed at the surface of donor cells could explain why some recipients’ immune systems mount an attack against the donor organ, while other patients tolerate the transplant for many years, when given similar immunosuppressive regimens. If the results of this study are confirmed in additional independent transplant cohorts (renal transplants, solid or hematologic transplants), they may prompt the design of prospective clinical trials to evaluate whether allocating organs to recipients with a combination of low allogenomics mismatch scores and low HLA mismatch scores improves long term graft outcome. A positive answer to this question could profoundly impact the current clinical and regulatory framework for assigning organs to ESRD patients.

In this study, we introduced the allogenomics concept to quantitatively estimate the histoincompatibility between an organ donor from an ABO compatible living donor and recipient outside of the HLA locus. We tested the simplest model derived from this concept to calculate an allogenomics mismatch score (AMS). We demonstrated that the AMS, which can be estimated before transplantation, helps predict post-transplantation kidney graft function. Interestingly, the strength of the correlation increases with the time post transplantation, an observation we make in both the discovery and validation cohorts. When testing the approach in a cohort of closely related pairs (transplantation mostly from related living donors), we observed that controlling for donor age at time of transplant was necessary to validate the association between eGFR and AMS. Donor age at time of transplant is a well-documented predictor of long-term graft function *(15)*. An alternative model where we control for donor age and time post-transplantation found a consistent effect of AMS across all three cohorts.

We chose to focus this study on living ABO compatible (either related or non-related) donors because kidney transplantation can be planned in advance and because differences in cold ischemia times and other covariates common in deceased donor transplants are negligible when focusing on living donors, especially in a small cohort.

The selection criteria for cadaveric donors include consideration of HLA matching and of the age of the donor. Compared to related donors we expect that the range of the AMS will be comparable to that in the discovery cohort (composed in majority of unrelated donors), Therefore, we expect that the allogenomics approach would also be predictive in cadaveric deceased donors, given that the AMS score should be high in these pairs. However, since many factors can independently influence graft function after transplantation from a cadaveric donor (e.g. cold ischemia time), potentially much larger cohorts may be required in such settings to achieve sufficient power to adequately control for the covariates relevant to cadaveric donors and to detect the allogenomics effect.

While several case-control studies have been conducted with large organ transplant cohorts, the identification of genotype/phenotype associations has been limited to the discoveries of polymorphisms with small effect, that have been reviewed in *(16)*, which have often not been replicated *(17–19)*. Such studies have observed small average effects measured across groups of transplants when our study is measuring moderate effects in individual transplants. Rather than focusing on specific genomic sites, the allogenomics concept sums contributions of many mismatches that can impact protein sequence and structure and could yield an immune response in the recipient.

While we have not attempted to optimize the set of sites considered to estimate the allogenomics mismatch score, it is possible that reduced and more focused subsets could increase the predictive ability of the score. For instance, the AMS could be applied to look for clusters of allogenomic mismatch sites in genes outside of the HLA loci. Such studies would require larger cohorts and may enable the discovery of loci enriched in allogenomics mismatches responsible for a part of the recipient alloresponse against yet unsuspected donor antigens. Their discovery might foster the development of new immunosuppressive agents targeting the expression of these immuno-dominant epitopes.

On the other hand, it is also possible that most polymorphisms that contribute to the score have a low frequency in the population (e.g., minor allele frequency less than 5%), which would make the identification of common sites of mismatches unlikely.

In this study, we propose several linear models that use the AMS to predict graft function after transplantation. A model that controls for donor age and time post transplantation, and includes the AMS, achieves an r2 of 0.22 and a RMSE of 15.49 when predicting MDRD eGFR. This model makes it possible to envision predicting graft function at an arbitrary future time after transplantation using genomic information available before transplantation. However, several multi-center independent validations will be essential to establish if prospective clinical trials are warranted. We recommend focusing such validation efforts on transplant pairs that are not familial, where the AMS effect appears to be maximal. We distribute the software that we developed to estimate the allogenomics mismatch score to facilitate further studies by others (see http://allogenomics.campagnelab.org).

## Materials and Methods

The study was reviewed and approved by the Weill Cornell Medical College Institutional Review Board (protocol #1407015307 “Predicting Long-Term Function of Kidney Allograft by Allogenomics Score”, approved 09/09/2014). The second study made on the French cohort was approved by the Comité de Protection des Personnes (CPP), Ile de France 5, (02/09/2014). Codes were used to ensure donor and recipient anonymity. All subjects gave written informed consent. Living donor ABO compatible kidney transplantations were performed according to common immunological rules for kidney transplantation with a mandatory negative IgG T-cell and B-cell complement-dependent cytotoxicity cross-match.

### Whole exome sequencing and genotyping

Briefly, genotypes of donors and recipients were assayed with exome sequencing (Illumina TruSeq enrichment kit for the Discovery Cohort and Agilent Haloplex kit for the Validation cohort and the French cohort. Reads were aligned to the human genome with the Last *(9)* aligner integrated as a plugin in GobyWeb *(8)*. Genotype calls were made with Goby *(10)* and GobyWeb *(8)*. Prediction of polymorphism impact on the protein sequence were performed with the Variant Effect Predictor *(20)*. Genes that contain at least one transmembrane segment were identified using Ensembl Biomart *(21)*.

### Discovery cohort: Transplant recipients and DNA samples

We selected 10 kidney transplant recipients from those who had consented to participate in the Clinical Trials in Organ Transplantation-04 (CTOT-04), a multicenter observational study of noninvasive diagnosis of renal allograft rejection by urinary cell mRNA profiling. We included only the recipients who had a living donor transplant and along with their donors, had provided informed consent for the use of their stored biological specimens for future research. Their demographic and clinical information is shown in Table 1. DNA was extracted from stored peripheral blood using the EZ1 DNA blood kit (Qiagen^®^) based on the manufacturer’s recommendation.

### Discovery cohort: Whole exome sequencing

DNA was enriched for exome regions with the TruSeq exome enrichment kit v3. Sequencing libraries were constructed using the Illumina TruSeq kit DNA sample preparation kit. Briefly, 1.8 μg of genomic DNA was sheared to average fragment size of 200 bp using the Covaris E220 (Covaris, Woburn, MA, USA). Fragments were purified using AmpPureXP beads (Beckman Coulter, Brae, CA, USA) to remove small products (<100 bp), yielding 1 μg of material that was end-polished, A-tailed and adapter ligated according to the manufacturer’s protocol. Libraries were subjected to minimal PCR cycling and quantified using the Agilent High Sensitivity DNA assay (Agilent, Santa Clara, CA, USA). Libraries were combined into pools of six for solution phase hybridization using the Illumina (Illumina, San Diego, CA, USA) TruSeq Exome Enrichment Kit. Captured libraries were assessed for both quality and yield using the Agilent High Sensitivity DNA assay Library Quantification Kit. Sequencing was performed with six samples per lane using the Illumina HiSeq 2000 sequencer and version 2 of the sequencing-by-synthesis reagents to generate 100 bp single-end reads (1×100SE).

### Validation cohort: Transplant recipients and DNA samples

We studied 24 kidney transplant recipients who had a living donor transplant at the NewYork-Presbyterian Weill Cornell Medical Center. This was an independent cohort and none of the recipients had participated in the CTOT-04 trial. Recipients were selected randomly based on the availability of archived paired recipient-donor DNA specimens obtained at the time of transplantation at our Immunogenetics and Transplantation Laboratory. The Institutional Review Board at Cornell approved the study. DNA extraction from peripheral blood was done using the EZ1 DNA blood kit (Qiagen®) based on the manufacturer’s recommendation.

### French cohort: Transplant recipients and DNA samples

We studied 19 kidney transplant recipients who had a living donor transplant at Tenon Hospital. This represented a third independent cohort. Recipients were selected randomly based on the availability of archived paired recipient-donor DNA specimens obtained either at the Laboratoire d'histocompatibilité, Hôpital Saint Louis APHP, Paris or during patient’s follow-up between October 2014 and January 2015. DNA extraction from peripheral blood was done using the Nucleospin blood L kit (Macherey-Nagel®) based on the manufacturer’s recommendation.

### Validation and French cohorts: Whole exome sequencing

The Validation and French cohorts were both assayed with the Agilent Haloplex exome sequencing assay. The Haloplex assay enriches 37 Mb of coding sequence in the human genome and was selected for the validation cohort because it provides a strong and consistent exome enrichment efficiency for regions of the genome most likely to contribute to the allogenomics contributions in protein sequences. In contrast, the TrueSeq assay (used for the Discovery Cohort) enriches 63Mb of sequence and includes regions in untranslated regions (5’ and 3’ UTRs), which do not contribute to allogenomics scores and therefore do not need to be sequenced to estimate the score. Libraries were prepared as per the Agilent recommended protocol. Sequencing was performed on an Illumina 2500 sequencer with the 100bp paired-end protocol recommended by Agilent for the Haloplex assay. Libraries were multiplexed 6 per lane to yield approximately 30 million PE reads per sample.

### Minor Allele Frequencies of the AMS Sites

We determined the minor allele frequency of sites used in the calculation of the allogenomics mismatch score using data from the NHLBI Exome Sequencing Project (ESP) release ESP6500SI-V2. We downloaded the data file ESP6500SI-V2-SSA137.protein-hgvsupdate.snps_indels.txt.tar.gz and extracted MAF in the European American population (EA) and in the African American population (AA) *(22)*. The ESP measured genotypes in a population of 6,503 individuals across the EA and AA populations using an exome-sequencing assay*(22)*. This resource made it possible to estimate MAF for most of the variations that are observed in the subjects included in our discovery and validation cohort.

### Overlap with EVP variants

Of 12,457 sites measured in the validation cohort with an allogenomics contribution strictly larger than zero (48 exomes, sites with contributions across 24 clinical pairs of transplants), 9,765 (78%) have also been reported in EVP (6,503 exomes).

### Sequence Data Analysis

Illumina sequence base calling was performed in the Weill Cornell Genomics Core Facility. Sequence data in FASTQ format were converted to the compact-reads format using the Goby framework [14]. Compact-reads were uploaded to the GobyWeb*(8)* system and aligned to the 1000 genome reference build for the human genome (corresponding to hg19, released in February 2009) using the Last *(9, 23)* aligner (parallelized in a GobyWeb *(8)* plugin). Single nucleotide polymorphisms (SNPs) and small indels genotype were called using GobyWeb with the Goby *(24)* discover-sequence-variants mode (parameters: minimum variation support=3, minimum number of distinct read indices=3) and annotated using the Variant Effect Predictor *(20)* (VEP version 75-75.7) from Ensembl. The data were downloaded as a Variant Calling format *(25)* (VCF) file from GobyWeb *(8)* and further processed with the allogenomics scoring tool (see http://allogenomics.campagnelab.org).

### Estimation of the Allogenomics Mismatch Score (AMS)

The allogenomics mismatch score Δ(*r,d*) is estimated for a recipient *r* and donor *d* as the sum of score mismatch contributions (see Fig. 1C, Equation 1).

**Equation 1** (reproduced from Fig. 1C).

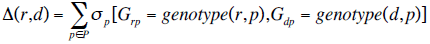

Contributions are observed for each polymorphic site *p* in a set *P*, where *P* is determined by the genotyping assay and analysis methods, and can be further restricted (*e.g*., to polymorphisms within genes that code for membrane proteins). Score mismatch contributions σ_p_(G_rp_,G_dp_) are calculated using the recipient genotype *G_rp_* and the donor genotype *G_dp_* at the polymorphic site *p*. Here, we consider that a genotype can be represented as a set of alleles that were called in a given genome. For instance, if a subject has two alleles at one polymorphic site, and we denote each allele A or B, the genotype at *p* is represented by the set {A,B}. This representation is general and sufficient to process polymorphic sites with single nucleotide polymorphisms or insertion/deletions.

Equation 2 describes how the individual score mismatch contributions are calculated at a polymorphic site of interest.

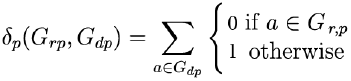

**Equation 2** (reproduced from Fig. 1C).

A contribution of 1 is added to the score for each polymorphic site where the donor genome has an allele (*a_dp_*) that is not also present in the recipient genome. When both donor and recipient genome are called at polymorphic site P, no contribution is added. For example, assuming a genomic site where the donor genome has two alleles, *i.e.*, *G_dp_*={A,B}, and the recipient genome is homozygote with *G_rp_*={A}. In this case, (*G_rp_*,*G_dp_*)=1. Fig. 1B presents additional examples of donor and recipient genotypes and indicates the resulting score contribution (the subscript *p* is omitted for conciseness). Score contributions are summed across all polymorphism sites in the set *P* to yield the allogenomic mismatch score (see Fig. 1C Equation 1).

### Selection of Informative Polymorphisms

The selection of the set of polymorphic sites *P* is important to the effectiveness of the approach. In the current method, we select exonic polymorphic sites that are (1) predicted to create non-synonymous change in a protein sequence, (2) are located in a gene that codes for one or more membrane proteins (defined as any protein with at least one predicted transmembrane segment, information obtained from Biomart *(21)*, Ensembl database 75). Additional filters can be applied to restrict *P*, which may lead to improved prediction of transplant clinical endpoints. Constructing additional filters will require the study of a larger training set of matched recipient and donor genotypes, which currently does not exist. It is possible that such study will indicate that other criteria than (2) also lead to predictive scores.

### Implementation: the Allogenomics Scoring Tool

We developed the *allogenomics scoring tool* to process genotypes in the VCF format and produce allogenomics mismatch scores for specific pairs of genomes in the input file. The *allogenomics scoring tool* was implemented in Java with the Goby framework and is designed to read VCF files produced by Goby and GobyWeb. The source code of the allogenomics scoring tools is distributed for academic and non-commercial purposes at http://allogenomics.campagnelab.org. The following command line arguments were used to generate the estimates described in this manuscript and can be run from the Allogenomics_Package file provided in supplementary. The genotype input file(s) necessary to reproduce these results (GobyWeb tags: JEOHQUR (2.3GB), YOOLWXH (83MB)) cannot be distributed through dbGAP (http://www.ncbi.nlm.nih.gov/gap), or an equivalent archive, because the consent form signed by the CTOT-04 participants is not compatible with such distribution of the subject information. A copy of the VCF files can be provided to the editors of the journal upon request should they wish to make it available to the reviewers upon condition of confidentiality during peer-review.

Pre-requisite to running the command lines: (1) You must have the Java runtime environment installed on your computer (the software has been tested with version 1.6) (2) You must define the environment variable ALLO to the location where you have downloaded the distribution of the allogemomics scoring tool (3). You must obtain the input VCF files and place them under: ${ALLO}/VCF_files_input/JEOHQUR-stats.vcf.gz ${ALLO}/VCF_files_input/YOOLWXH stats.vcf.gz.

<u>Estimating allogenomics mismatch scores on the Discovery cohort:</u>

java -Xmx4g -jar allogenomics-1.1.7-scoring-tool.jar \

           --input ${ALLO}/VCF_files_input/JEOHQUR-stats.vcf.gz \

           -p ${ALLO}/Pair_files/Discovery_cohort.pairs.tsv \

           -a                          Annotation_files/All_protein_coding_Ensembl_75.gtf \

           --output ${ALLO}/Output/TM-Discovery.tsv \

           --output-format TSV --only-non-synonymous-coding --vep \

           --consider-indels --minimum-depth 10 --max-depth 500 \

           -t ${ALLO}/Annotation_files/TrM-Transcript_Ensembl_75.tsv \

           --clinical

<u>Estimating allogenomics mismatch scores on the Validation cohort:</u>

java -Xmx4g -jar allogenomics-1.1.7-scoring-tool.jar \

           --input ${ALLO}/VCF_files_input/JEOHQUR-stats.vcf.gz \

           -p ${ALLO}/Pair_files/Validation_cohort.pairs.tsv \

           -a ${ALLO}/Annotation_files/All_protein_coding_Ensembl_75.gtf \

           --output ${ALLO}/Output/TM-Validation.tsv \

           --output-format TSV --only-non-synonymous-coding --vep \

           --consider-indels --minimum-depth 10 --max-depth 500 \

           -t ${ALLO}/Annotation_files/TrM-Transcript_Ensembl_75.tsv \

           --clinical --measured-sites SitesHaloplexExome.tsv

<u>Estimating allogenomics mismatch scores on the French cohort:</u>

java -Xmx4g -jar allogenomics-1.1.7-scoring-tool.jar \

           --input ${ALLO}/VCF_files_input/YOOLWXH-stats.vcf.gz \

           -p Pair_files/French_cohort.pairs.tsv \

           -a ${ALLO}/Annotation_files/All_protein_coding_Ensembl_75.gtf \

           --output ${ALLO}/Output/TM-French_cohort.tsv \

           --output-format TSV --only-non-synonymous-coding \

           --vep --consider-indels --minimum-depth 10 \

           --max-depth 500 \

           -t ${ALLO}/Annotation_files/TrM-Transcript_Ensembl_75.tsv --clinical --no-dash

<u>Estimating allogenomics mismatch scores on merged discovery and validation cohorts:</u>

java -Xmx4g -jar allogenomics-1.1.7-scoring-tool.jar \

           --input ${ALLO}/VCF_files_input/JEOHQUR-stats.vcf.gz\

           -p Pair_files/Discovery+Validation_cohort.pairs.tsv \

           -a ${ALLO}/Annotation_files/All_protein_coding_Ensembl_75.gtf \

           --output ${ALLO}/Output/TM-Discovery+Validation.tsv \

           --output-format TSV --only-non-synonymous-coding \

           --vep --consider-indels --minimum-depth 10 \

           --max-depth 500 \

           -t ${ALLO}/Annotation_files/TrM-Transcript_Ensembl_75.tsv --clinical

Estimating allogenomics mismatch score limited to Illumina GeneChip660W loci on the validation cohort:

java -Xmx4g -jar allogenomics-1.1.7-scoring-tool.jar \

           --input ${ALLO}/VCF_files_input/JEOHQUR-stats.vcf.gz \

           -p ${ALLO}/Pair_files/Validation_cohort.pairs.tsv \

           -a ${ALLO}/Annotation_for_660W/Human660W_Gene_Annotation_hg19-ilmn.tsv \

           --output ${ALLO}/Output/TM-Validation_Illumina660W.tsv \

           --output-format TSV --only-non-synonymous-coding --vep \

           --consider-indels --minimum-depth 10 --max-depth 500 \

           -t ${ALLO}/Annotation_for_660W/TM-as-gene-names_for_Illumina660W.tsv \

           --clinical --measured-sites sites-660W.tsv

### Statistical Analyses

Analyses were conducted with either JMP Pro version 11 (SAS Inc.) or metaR (http://metaR.campagnelab.org). Figures 2, 3 and 4, as well as SF1B, SF1C, SF2B, SF3C were constructed with metaR analysis scripts and edited with Illustrator CS6 to increase some font sizes or adjust the text of some axis labels. The model that includes the time post-transplantation as a covariate was constructed in metaR and JMP. The R implementation of train linear model uses the lm R function. This model was executed using the R language 3.1.3 (2015-03-09) packaged in the docker image fac2003/rocker-metar:1.4.0 (https://hub.docker.com/r/fac2003/rocker-metar/). Models with random effects were estimated with metaR 1.5.1 and R (train mixed model and compare mixed models statements, which use the lme4 R package*(26)*). Comparison of fit for models with random effects was obtained by training each model alternative with REML=FALSE an performing an anova test, as described in *(27)*.

## Acknowledgments

We thank Dr. Joseph Schwartz and Dr. Samprit Banerjee for independent critical review of the manuscript. **Funding**: This study was funded from institutional funds (MS and FC). **Author contributions**: Study design and manuscript preparation (LM, TM, MS, FC). Statistical analyses (LM, JRL, FC). Bioinformatics analyses (LM, FC). Wrote the allogenomics scoring tool (FC). Genomics Core, Exome Sequencing (HS, JX). Collating clinical data from hospital systems for CTOT-04 and validation study (TM, MB, CL). HLA Typing, validation cohort (DD), French cohort (CS, MC). Samples and clinical information for Discovery cohort (JJF, MMA). French cohort: samples (LM, NO, ER), clinical information (LM, NO, ER). **Competing interests:** Laurent Mesnard, Thangamani Muthukumar, Manikkam Suthanthiran and Fabien Campagne disclose that they are named inventors in a filed international patent application entitled “A METHOD TO MATCH ORGAN DONORS TO RECIPIENTS FOR TRANSPLANTATION”. **Data and materials availability**: de-identified data and analysis software are distributed online. A copy of the genotype files can be provided to the journal upon request should the editors wish to make them available to the reviewers upon condition of confidentiality during peer-review.

## Supplementary Materials

Supplementary materials and methods appendix containing:

**Fig. S1**. Model trained on the Discovery cohort applied to the Validation cohort

**Fig. S2**. Model trained on the Validation cohort applied to the Discovery cohort

**Fig. S3**. Effect of genotyping platform on future replication studies.

**Supplementary Figures**

**Figure S1.**
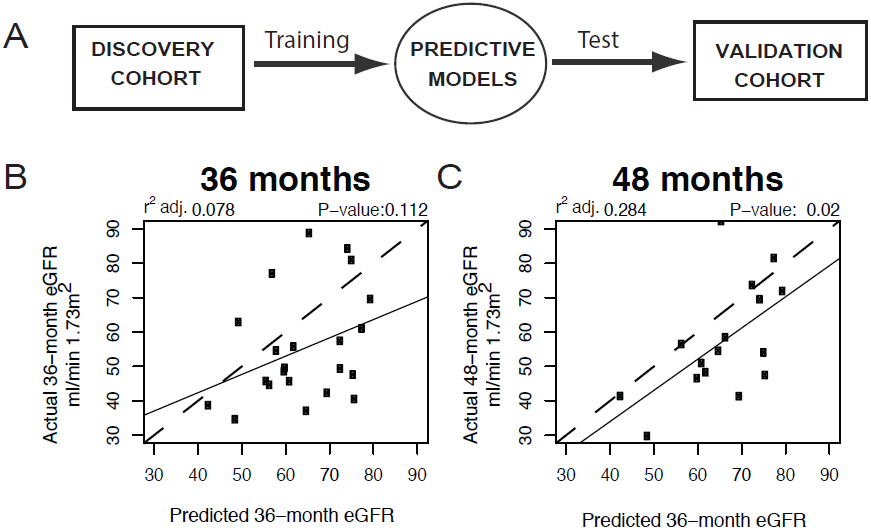
Model trained on the Discovery cohort applied to the Validation cohort. A) We trained a model to predict eGFR on the discovery cohort (using eGFR at 36 months) and used the trained, fixed, model to predict eGFR at 36 months and 48 months for recipients of the Validation cohort. The trained model was eGFR= 107.39547- -0.03974*AMS. Correlation between predicted eGFR and observed eGFR on the Validation cohort at 36 (B) and 48 (C) months post transplantation. Dashed lines indicate the diagonal and solid lines the regression lines.

**Figure S2.**
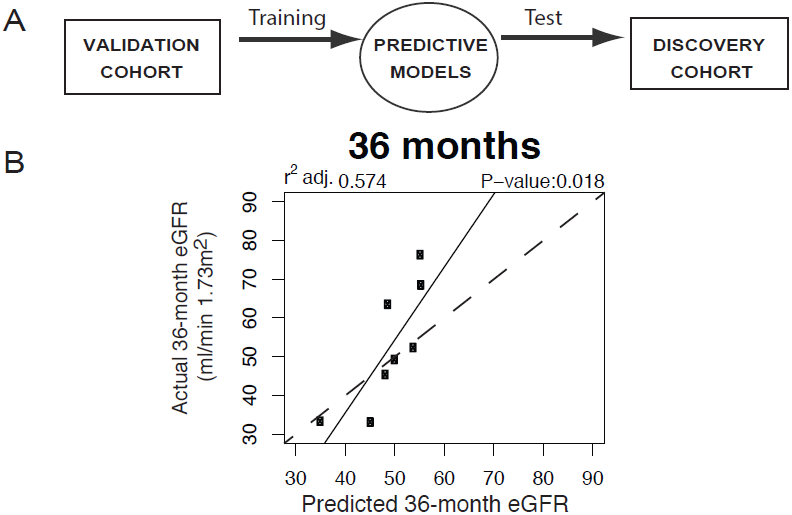
Model trained on the Validation cohort applied to the Discovery cohort. A) We trained models to predict serum creatinine and eGFR on the validation cohort and used the trained, fixed, model to predict serum creatinine and eGFR for recipients of the Discovery cohort. B) Correlation between the eGFR predicted by the fixed model and that observed in the Discovery cohort. The trained model was eGFR= 78.20459 -0.02114*AMS. Dashed line indicates the diagonal and solid line the regression lines.

**Figure S3.**
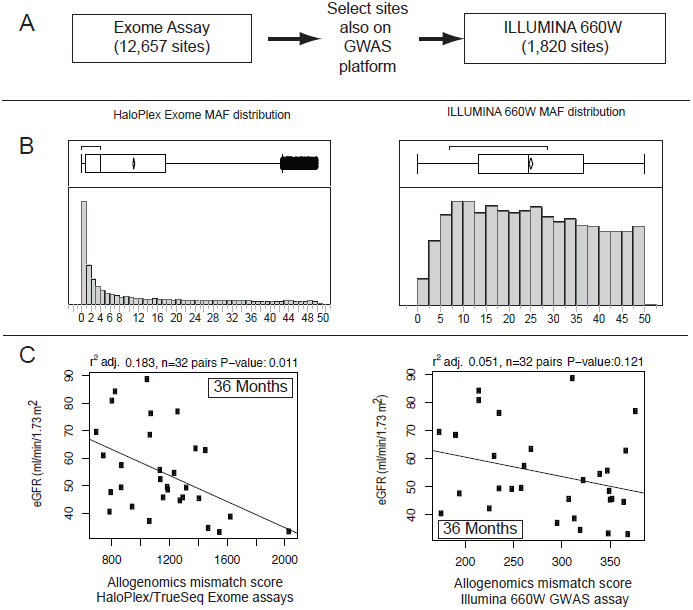
Effect of genotyping platform on future replication studies. In this analysis, we estimate how well the allogenomics mismatch score could be evaluated with the genotyping array technology frequently used in GWAS studies. Analyses are done on the combined Discovery and Validation cohorts (n=32 pairs with 36-month eGFR, 64 exomes). A) The allogenomics mismatch score evaluated with the Illumina TrueSeq or Agilent Haloplex exome platforms takes advantage of 12,657 genomic sites to estimate allogenomic contributions in transmembrane proteins. Only sites where an allogenomics mismatch score contribution different from zero are counted. We filtered the exome genomic sites to exclude sites not found on the Illumina 660W genotyping platform (used in *(13)*). After filtering, the allogenomics score is estimated with 1,820 remaining genomic sites. B) The minor allele frequency (MAF) of the alleles described at each set of genomic sites is shown as a histogram (MAF is estimated from the EVP database, see Methods). Exome sequencing is an assay that directly observes variations in an individual DNA sample. The MAF distributions confirm that exome sequencing helps estimate contributions from many rare (MAF<5%) polymorphisms, whereas the chip genotyping platform estimates the score based on contributions from frequent alleles. C) The scatterplot of the relationship between 36-month eGFR and the score estimated from the exome sites, or the subset of sites also measured by the GWAS platform. While some trend is still visible with sites measured on the GWAS platform, more samples would be needed to reach significance in the combined Discovery and Validation cohorts (n=34 pairs). Note that the magnitude of the scores is smaller on the GWAS platform because fewer contributions are summed. In contrast, the exome assays (Illumina TrueSeq for the Discovery cohort or Agilent Haloplex for the Validation cohort) result in stronger and significant correlations in the same set of samples.

